# Distinct tumor-immune ecologies in NSCLC patients predict progression and define a clinical biomarker of therapy response

**DOI:** 10.1101/2022.10.22.513219

**Authors:** Sandhya Prabhakaran, Chandler Gatenbee, Mark Robertson-Tessi, Jhanelle E. Gray, Scott Antonia, Amer A. Beg, Robert A. Gatenby, Alexander R. A. Anderson

## Abstract

We investigated multiplexed histological images from nine patients (both pre- and on-treatment) with immunotherapy-refractory Non-Small Cell Lung Cancer (NSCLC) treated with an oral HDAC inhibitor (vorinostat) combined with a PD-1 inhibitor (pembrolizumab). Patient responses comprised of either stable disease (SD) or progressive disease (PD). We built an extensive multiplexed-image analysis pipeline involving both cell segmentation and quadrats, coupled with spatial statistics, machine learning, and deep learning to analyze the spatial and temporal features that predict disease progression and identify potential clinical biomarkers. We found that distinct spatial immune ecologies exist between SD and PD patients. We also demonstrate that tumors from PD patients are already characterized by an immune-suppressive environment prior to treatment. Finally, we show that the learned spatial ecologies can predict disease progression better than PD-L1 status alone, suggesting these ecologies can be used as potential companion biomarkers with PD-L1 in NSCLC. These findings will be investigated in a larger-cohort study generated from an ongoing clinical trial (NCT02638090) that includes a wider range of responses including complete and partial responders. Additionally, the computational infrastructure developed in this study can be generalized to any cancer type.

## Introduction

Despite major advances in elucidating the molecular mechanisms of lung cancer, and improvements in prognosis, diagnosis and treatment, lung cancer remains the leading cause of cancer death worldwide^1^. Non-Small Cell Lung Cancer (NSCLC) accounts for about 85% of all cases^2^. Disease progression and treatment response in NSCLC vary widely among patients. As targeted molecular therapies, immunotherapies, and combination therapies have become central in managing patients with NSCLC, the ability to identify which patients are appropriate for each therapeutic approach has become even more important. This has accelerated the research to develop *biomarkers* for identifying patients who would likely respond to therapy. Additionally, biomarkers monitor disease development and help predict disease outcomes and treatment response^3,4^. In the case of immunotherapy, existing treatment response biomarkers (PD-L1) are useful, but there are well recognized limitations^3^, such as, the spatial and temporal heterogeneity of PD-L1 expression, the dynamic nature of PD-L1 expression, and different methods used to stain and measure PD-L1 expression within tumors. These may account for the challenges in correlating PD-L1 expression with patient response^5–8^. Therefore, identifying novel biomarkers to accurately predict the likelihood of response to immunotherapy is crucial in treatment selection and planning for each NSCLC patient.

Multiplexed imaging of tissues, and their analysis, is an emerging and proficient approach aiding early detection, clinical cancer diagnosis, treatment planning, and prognosis^9–13^. These images enable the precise determination of spatial distributions of cells and cellular states and the characterization of tumor-immune interactions *in situ* at the single-cell level. Furthermore, they allow simultaneous detection of various protein biomarkers on the same tissue sample, permitting molecular and immune profiling of NSCLC, while preserving tumor architecture^10,14^, and enable the direct quantification of response to a given treatment. However, quantitative image analytic tools must be developed to enable these data to drive predictive biomarkers of response.

The analysis of multiplexed images generally utilizes one of the two state-of-the-art image analysis approaches, i.e., cell segmentation and quadrats. Single-cell data, generated using cell segmentation methods^15^, is a fundamental step in many biomedical studies. It quantifies the marker expression of individual cells in a field of view (FoV) and is used to identify cell morphologies, cell subpopulations and neighborhoods, and the stochasticity of marker expression. The approach can be used to extract tumor, stromal, and invasive regions, as well as identify the cells that colocalize in such regions. Dimensionality reduction of the single-cell data can aid with visualization of this high-dimensional data. Single-cell data relies on accurate cell segmentation which can be cumbersome depending on the number of samples and the technical noise of the samples.

The second approach, quadrat analysis, is an important process in image analysis where the high-dimensional image is sequentially tiled, summarized, and classified. Quadrats can be used to identify aggregate marker expression and differential markers across the population of cells in a sample and is typically used to detect interesting regions across high-dimensional images. This does not require any cell-specific information, making it a simple yet effective approach. In a significant amount of high-throughput image analysis research, using tiles for pattern matching and classification is standard^15^. As ecologists have long worked with quadrat count data, an additional benefit of tiling the images is that it allows one to leverage many of the spatial analyses originating from ecology, an approach taken in several studies of the tumor microenvironment (TME)^16–21^.

With recent advances in spatial omics, there has been an increased interest in exploring spatial tumor ecologies, i.e., the cellular and quadrat patterns and interactions within the tumor^22–24^, how these tumor ecologies are related to patient response to treatment, and how these are impacted by single-agent drugs or immune checkpoint inhibitor (ICI) combination therapies. As tumors are dynamic, rapidly evolving and undergoing ecological transformations, the study of cancer evolution has to incorporate an understanding of the dynamic ecology to explain heterogeneity, behavior, and eco-evolutionary feedbacks^25–27^.

In this study, our goal is to uncover the evolving spatiotemporal ecologies that drive progression in NSCLC patients and identify predictive biomarkers of response, by leveraging multiplexed images. Here, we analyzed multiplex histological images from pre- and on-treatment (days 15-21) in nine patients with immunotherapy-refractory NSCLC treated with an oral HDAC inhibitor (vorinostat) combined with a PD-1 inhibitor (pembrolizumab)^28^. We developed a computational multiplexed-image analysis pipeline using both cell segmentation and quadrats to better understand the spatial and temporal features of multiplexed NSCLC images, and their ability to predict disease progression, as well as identify clinical biomarkers. Using cell segments and quadrats, we defined distinct spatial neighborhoods and associated cellular interactions, and inferred spatial ecologies using only patient-level labels. An important aspect of our study is that combining both approaches allowed us to quantify the unique spatial ecologies in stable disease (SD) and progressive disease (PD) patients, and enabled response predictions based on the abundances and diversity of discovered marker phenotypes and neighborhoods. These approaches, if validated in larger cohorts, may provide improved predictive histological biomarkers from pretreatment samples to drive subsequent treatment decisions. Further, these approaches can be generalized to study multiplexed images from any cancer type.

## Results

### Study design and data source

We have used 92 PerkinElmer Vectra Field of Views (FoVs) (hereby referred to as FoVdata) from nine patients with immunotherapy-refractory NSCLC with progression^28^. They were part of a randomized Phase I trial (NCT02638090) and were treated with an oral HDAC inhibitor (vorinostat) combined with a PD-1 inhibitor (pembrolizumab). Tumor biopsies were collected from all patients both pre- and on-treatment (days 15-21). Patient responses were classified into two groups: those who remained in stable disease (SD) after 24 weeks; and those who had progressive disease (PD) before 24 weeks^28^. Of the nine patients, five qualified as SD and four as PD. There are 58 images from the SD patients and 34 images from the PD patients.

Using the seven-color multiplex fluorescence assay by PerkinElmer-Opal-Kit, each image is stained for seven markers: DAPI (nuclei); CD3 (T cells); CD8 (effector T cells); FoxP3 (regulatory T cells); PD-1 (inhibitory receptor on immune cells); PD-L1 (immune checkpoint marker); and Pan-cytokeratin (PanCK, epithelium). We had a median of 10 images per patient in FoVdata, split evenly across pre- and on-treatment, except for one patient for whom we had access to only a single image taken pre-treatment. We analyzed the FoVdata to identify spatial ecologies and to make disease progression predictions (**Figure 1**). Both stromal and tumor regions were imaged in the FoVdata. We summarize FoVdata in **Table 1**. An exemplar FoV with its corresponding cell segments and quadrats is shown in **Figure 2A**, along with its individual markers and composite marker image (**Appendix Figure 4**).

**Table 1.** Table summarizing FoVdata.

| NSCLC multiplexed images | #Patients | #Images | #Response | #Images per response | #Cells | #Quadrats 100$\mu\text{m}$ x 100$\mu\text{m}$ | Format | Dimension per image | Size per image |
| --- | --- | --- | --- | --- | --- | --- | --- | --- | --- |
| **FoVdata** | 9 | 92 FoVs | 5 SD (pre + on) | 58 FoVs | 80K<br>Pre: 42K<br>On: 38K | 3.5K<br>Pre: 2K<br>On: 1.5K | TIFF | 1400 x 1000 | 10-50 MB |
|  |  |  | 4 PD (pre + on) | 34 FoVs | 41K<br>Pre: 21K<br>On: 20K | 2K<br>Pre: 994<br>On: 1K |  |  |  |

**Figure 1:**
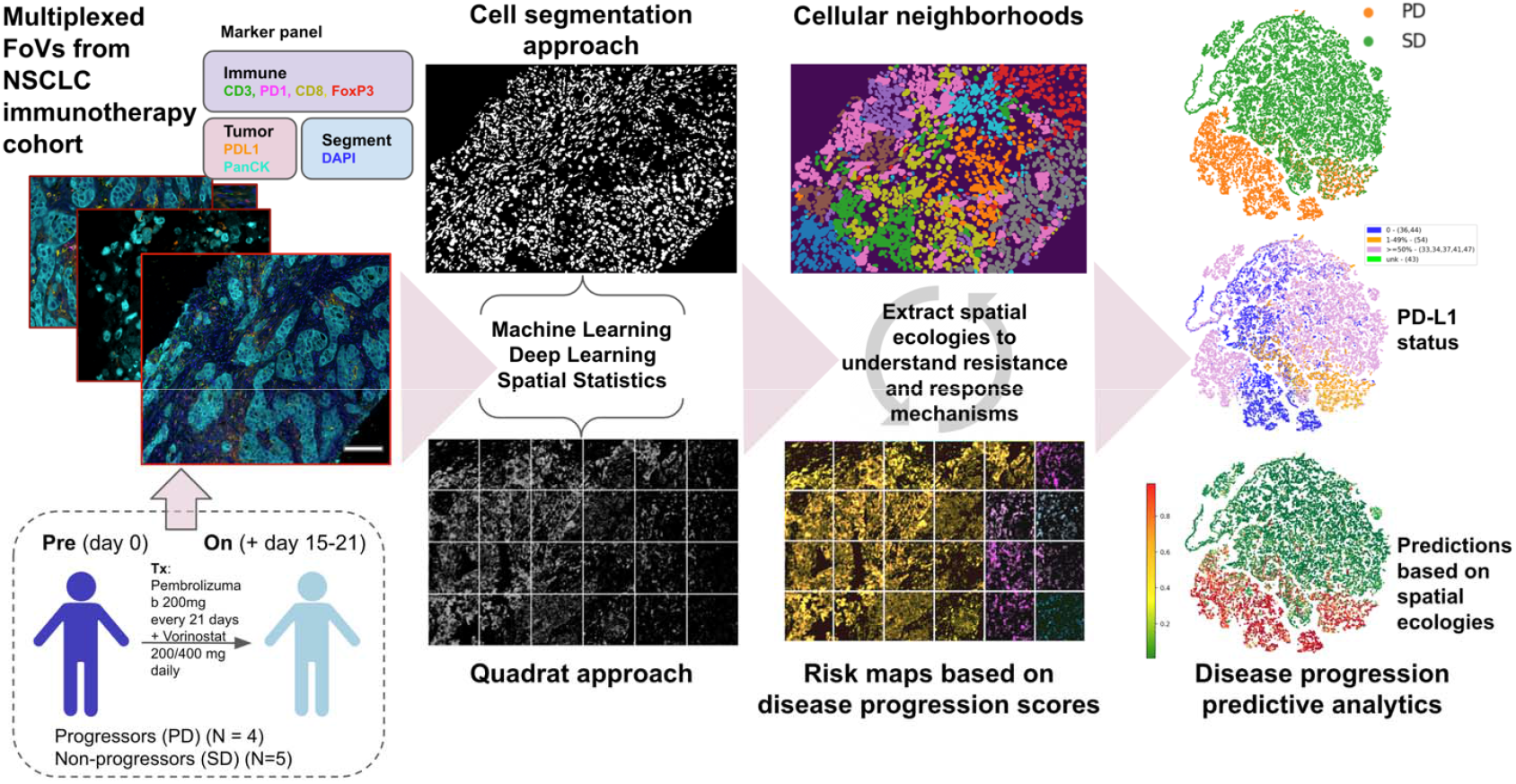
Graphical abstract showing the two computational approaches developed in this paper to study spatial and temporal interactions of tumor and immune cells using multiplexed images from Moffitt’s NSCLC immunotherapy trial cohort. A longitudinal cohort of nine NSCLC patients were imaged before (Pre) and during (On) treatment. For the single cell approach, we extract the cell segments per Field of View (FoV) using a U-Net, build a count matrix with cells as rows and markers as columns, and cluster the count matrix to identify spatially heterogeneous cell types. For the quadrat approach, we view each patient sample as a collection of fixed-size grids, called quadrats, to study the marker distribution and to detect global changes across the spatially diverse cell types. Using an array of downstream analyses involving spatial statistics, machine learning and deep learning, we identify key cellular neighborhoods (ecologies) distinguishing PD from SD patients and generate risk maps that aid actionable clinical decisions. (Scale bar = 200 pixels/µm). We build predictive models that leverage these ecologies to stratify the two response groups, and compare these predictions to the standard PD-L1 status-based predictions.

**Figure 2:**
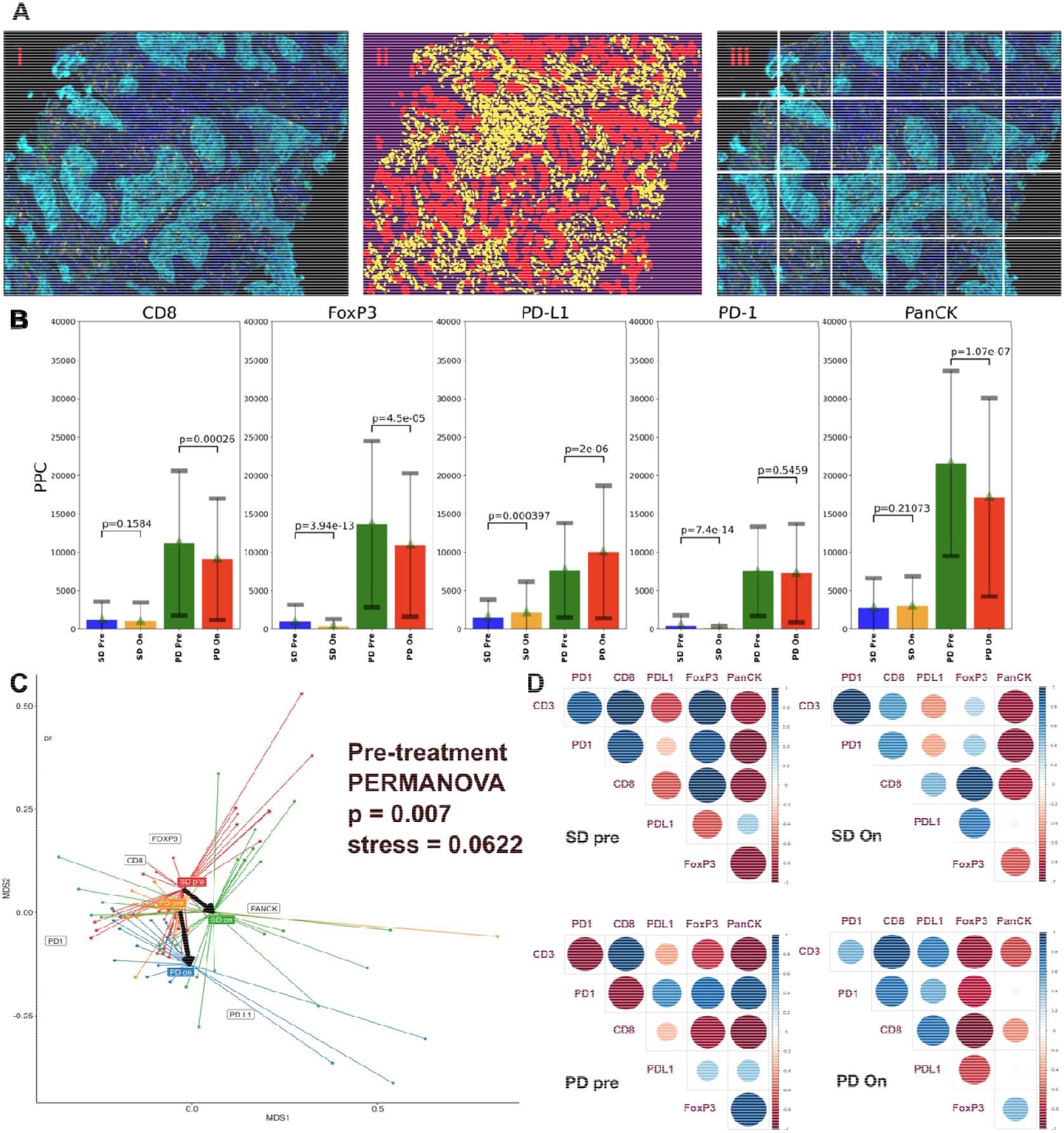
Non-spatial analyses suggest that the pre-treatment ecology may be predictive of response. **A**. (i) An example NSCLC FoV which is a 7-stain Vectra image with DAPI (blue; nuclei), CD3 (green; T cells), CD8 (yellow; effector T cells), FoXP3 (red; regulatory T cells), PD-1 (pink; inhibitory receptor on immune cells), PD-L (orange; immune checkpoint marker), and pan-cytokeratin (PanCK, cyan; epithelium). Corresponding image showing **(ii)** cell segments (tumor cells shown in red) and **(iii)** image quadrats. **B**. Bar plots showing marker abundances (based on average positive pixel counts (PPC)) across SD and PD patients, before and during treatment, for each marker. The Bonferroni-adjusted p-value with significance for markers, pre and during treatment, are shown; p <= 5.00e-02 is significant. **C**. The NMDS (non-metric multidimensional scaling) projection of the average positive pixel counts across all FoV quadrats. Shown also is the PERMANOVA test indicating that the centroids or spread of each pre-treatment grou are significantly different (p-value = 0.007). **D**. Correlation matrix heatmaps computed using quadrats across patient response groups showing pairwise marker correlations (blue indicates positive correlation and red negative correlation). Size of each circle corresponds to the magnitude of the pairwise correlation represented.

### Non-spatial analyses indicate PD and SD patients have significantly different ecologies

Through our non-spatial analysis using positive pixel counts (PPCs) derived from FoVdata quadrats, it is evident that PD tumors demonstrate increased marker levels throughout (pre- and during-treatment) (**Figure 2B**). We overlay the NMDS (non-metric multidimensional scaling) projection of the average PPCs across all FoV quadrats with the PERMANOVA test (see Methods) to depict that the centroids, indicating the spread of each group, are different; this reinforces that PD and SD have significantly different pre-treatment ecologies (p-value = 0.007). Since each groups’ centroid shifted while on treatment, this also indicates that treatment itself alters the tumor ecology, as shown by the arrows in **Figure 2C**. In **Figure 2D**, we plot the pairwise marker correlations across SD and PD patients, computed based on PPCs, before and during treatment, and again note that marker correlations vary across response groups. For instance, in PD patients (both pre- and during-treatment), the correlations between FoxP3 with PanCK, and PD-1 with PanCK, are noticeably higher (blue) than in SD patients (red). These significant differences in composition derived from basic non-spatial analyses suggest that the pre-treatment ecology may be sufficiently different in PD and SD to be predictive of response.

### Spatial analyses highlight distinct spatial communities in SD and PD patients

We further study the spatial heterogeneity of tumors across response groups, by investigating the differences pertaining to the spatial organization of the tumor with respect to its microenvironment in the FoVdata cohort using cell segmentation and quadrat approaches. By clustering cells from all FoVs along with their spatial neighbors, we obtain twelve distinct CNs (**Figure 3A, Appendix Figure 2A**). An example of the spatial orientation of the CNs mapped onto a FoV is shown in **Figure 3B**. The t-SNE rendering of these CNs demonstrates that there are characteristic cellular neighborhoods defining PD and SD patients (**Figure 3Ci, ii**) such as PanCK+FoxP3+PD-1+ for PD patients (clusters 6 and 11) and PanCK+PD-L1+ with CD3+CD8+ for SD patients (clusters 4, 5, 7 and 8). It is evident that prior to treatment, PD patients show a higher number of cells expressing FoxP3+PD-1+ and PanCK+ (**Appendix Figure 2B)**. On the other hand, there is always an increased number of cells expressing CD3+, CD8+, PD-L1+ and PanCK+, both before and during treatment in SD patients. These interactions have been further reinforced visually using an image t-SNE generated using Mistic^29^ (**Appendix Figure 3**). Further, the tumor boundary analyses using convex hulls indicates that: a) for both pre- and on-treatment patients, there is a higher presence of immune cells (primarily, FoxP3) within the tumor of PD patients, and more so within PD patients during treatment (**Appendix Figure 1iii** and **Appendix M.3**); b) a higher abundance of tumor and immune cells at the tumor boundary (**Appendix Figure 1iv**) in PD than SD patients; and c) the potential to identify groups of immunologically hot, warm or cold tumors across patients based on immune cell colocalization at the tumor margins and tumor regions (**Appendix Figure 3**). Together, these results support that there are distinct cellular compositions, both within the tumor regions and at the tumor boundaries, for PD and SD patients.

**Figure 3:**
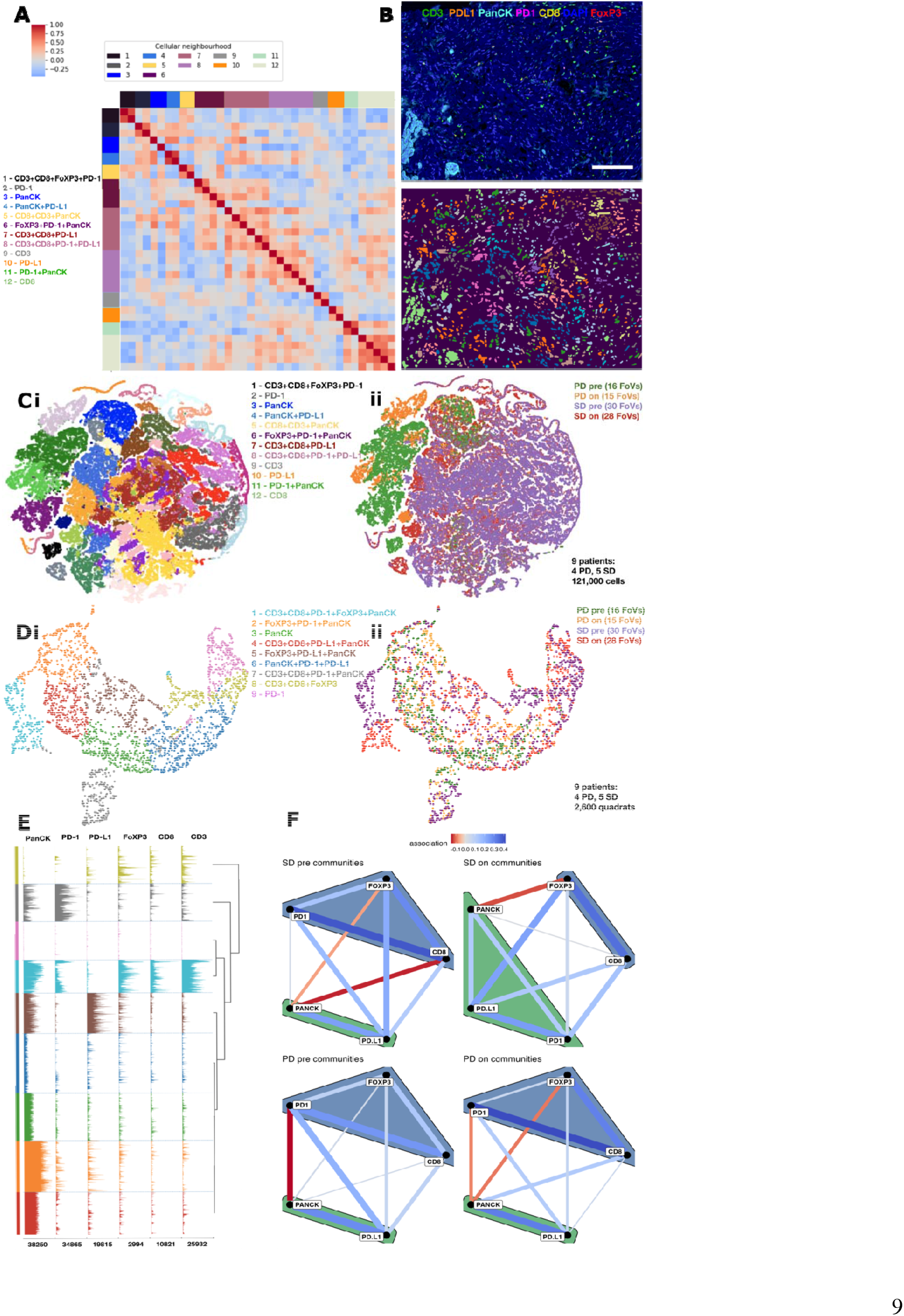
Diverse cellular and quadrat neighborhoods captured across patient categories. **A**. Heatmap showing 12 distinct cellular neighborhoods (CNs) based on six markers and their respective z-scored frequencies within each CN (pooled data across both SD and PD groups and across pre and on treatment). **B**. (**Top**) Exemplar multiplex FoV in a seven-color overlay image with DAPI (blue; nuclei), CD3 (green; T cells), CD8 (yellow; effector T cells), FoXP3 (red; regulatory T cells), PD-1 (pink; inhibitory receptor on immune cells), PD-L1 (orange; immune checkpoint marker), and pan-cytokeratin (PanCK, cyan; epithelium). Scale bar = 200 pixels/µm. (**Bottom**) FoV showing the twelve clustered CNs mapped. **C**. 2D t-SNE rendering of all the cells (~121,000) colored **(i)** by CNs, **(ii)** by patient category. **D**. 2D t-SNE rendering of all the quadrats (~2,600) colored **(i)** by QNs, **(ii)** by patient category. **E**. Traceplot showing marker contribution across each QN. Rows are QNs and columns are markers. Numbers at the base of each column indicate total number of cells across all quadrats for that specific marker. Legend similar to D (i). **F**. Optimal network of spatial community associations obtained using a Gaussian copula graphical lasso (GCGL) shown per patient category. These associations are indicated by the green/blue polygons capturing sets of markers and are obtained using the Leiden clustering algorithm.

Next, we cluster the quadrats generated from FoVdata using a Gaussian mixture model (GMM)^30^ and illustrate the inferred quadrat neighborhood (QN) clusters on t-SNE renderings (**Figure 3D**) and the marker contributions per cluster (**Figure 3E**). It is again evident that the QNs capture the distinct ecologies defining PD and SD patients such as PanCK+FoxP3+PD-1 (cluster 2: orange; cluster 7: gray) for PD patients, and PanCK+PD-L1 with CD3+CD8 (cluster 1: sky blue; cluster 5: brown) for SD patients. We construct species association networks^31–33^ (SANs) (**Figure 3F**) using the Gaussian copula graphical lasso (GCGL)^34,35^ method to identify marker interactions. We find that SANs confirm: the pre-treatment association of FoxP3 with PD-1 and PanCK in PD patients; and the on-treatment association CD8 with PD-L1 and PanCK in SD patients. Since the Gaussian copula can directly model covariance and strength between markers, the GCGL creates a single association network with all direct links between markers and the strength of association increases from red (negative/dispersed) to blue (positive/clustered). Community associations are shown by the larger polygons capturing sets of markers that are clustered using the Leiden algorithm^36^. By collectively examining each marker’s importance, response, and interactions across quadrats, we again observe distinct cellular communities distinguishing PD and SD patients.

Through these clustering analyses, we demonstrate that PD tumors are characterized by an immune-suppressive environment (due to presence of FoxP3+ and PD-1+ cells) prior to treatment. PD-1+ immune cells may have other checkpoint receptors such as LAG3, TIM3 or TIGIT that would indicate a deeper immune cell exhaustion state^37^ further reducing the response to treatment. On the other hand, SD tumors show the presence of higher immune cell infiltration indicating durable clinical benefit^38^.

### Fundamentally distinct ecologies across PD and SD patients enable disease progression prediction and clinical biomarker identification

For the FoVdata cohort, we trained a support vector machine (SVM) using the single cell data to predict response to therapy at the patient level. Using the leave one out cross validation (LOOCV) approach, the SVM achieves an overall mean accuracy of classification of 87.5%, with class-specific mean accuracy of 100% (SD) and 66.7% (PD) (**Figure 4A row1)**. To identify the marker importance per patient and therefore for each response group, we assess the weights obtained from the SVM. These weights represent the vector coordinates which are orthogonal to the SVM hyperplane, and their direction indicates the predicted class. The absolute size of the coefficients in relation to each other can then be used to determine marker importance for the prediction task. The marker importance per response group, generated by the SVM, indicates that CD3+CD8+PD-L1 are important markers for SD patients and FoxP3+PD-1 for PD patients, respectively (**Figure 4B**).

**Figure 4:**
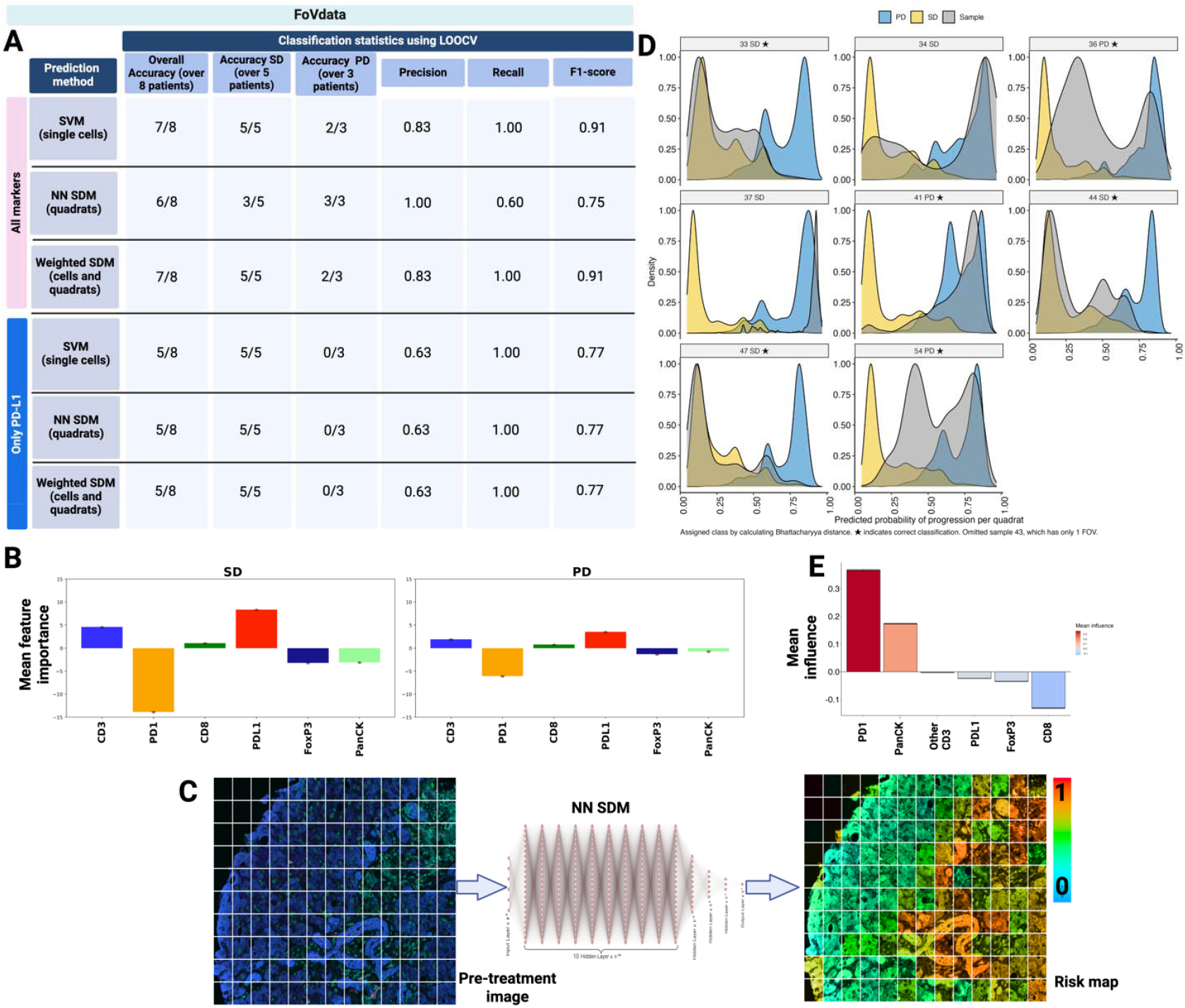
**A**. Disease progression prediction metrics using a Support Vector Machine (SVM) trained on cells, neural network species distribution model (NN SDM) trained on quadrats, and weighted SVM trained using quadrat probability distributions, from pre-treatment FoVdata images. Metrics shown for all markers (analyzed in their spatial context) versus PD-L1 alone. Metrics calculated based on the number of patients correctly classified. **B**. Mean marker importance per response group obtained by the SVM, where the absolute value is indicative of th effect each marker has on making accurate predictions. **C**. Disease progression predictions using a neural network species distribution network (NN SDM) trained on quadrats from pre-treatment images. The spatial co-localization of cells within each 50μm x 50μm quadrat (outlined in white) from pre-treatment biopsies (left) were used to train a NN SDM (middle) that predicts the probability a patient will progress while on treatment. The NN SDM prediction probabilities are projected back on to the image, potentially identifying regions driving disease progression (right). The images here are from one FoV from a patient that had progressive disease (PD). **D**. Using the LOOCV for the NN SDM, we show the distribution of quadrat probabilities for the test (sample) patient i comparison to the trained SD and PD quadrat probability distributions. A ‘*’ indicates correct classification. **E**. A feature-importance analysis using quadrats predictions from the NN SDM used to predict PD status. Values her thus indicate the importance each marker has in predicting whether the patient will progress while on treatment. Here, positive values indicate the marker is important in predicting PD, while negative values indicate importance in predicting SD.

A neural network (NN) species distribution model (SDM) (see Methods, **Appendix Figure 7)** trained on quadrat counts has an overall mean accuracy of 75% with class-specific mean accuracy of 60% (SD) and 100% (PD) (**Figure 4A row2**). The distribution of quadrat probabilities of PD classification, generated through the NN SDM (**Figure 4C**), could be used to assess risk of progression, as on average patients with stable disease (SD) had lower quadrat probabilities than PD patients. This distribution for each patient tested under the LOOCV method is compared to the distributions of both SD and PD patients generated during training, using the Bhattacharyya distance (BD)^39,40^ as the distance/similarity metric. The test distribution is assigned to either SD or PD based on the closest BD distance, i.e. the more similar distribution of progression probabilities (**Figure 4D**). A feature-importance analysis^41,42^ reveals that PD-1, FoxP3, CD8 and PanCK had the greatest impact on the NN SDM’s ability to accurately predict treatment response, suggesting these markers play the most important roles in determining whether a patient responds to treatment (**Figure 4E**). Positive values indicate the marker is important in predicting PD, while negative values indicate importance in predicting SD. Further, the NN SDM predictions allow us to create a *risk map* that identifies areas of the tumor associated with different probabilities of overall patient progression while on treatment (**Figure 4C**).

In addition to the above, we perform three crossover experiments involving cell segments and quadrats. In the first crossover experiment, we implement a weighted SVM. For this, each cell’s marker expression is weighted by the average of the corresponding quadrat progression probabilities (as obtained from the NN SDM), and this updated matrix is used to train the SVM. With this approach, while the predictions at the patient level remain the same (**Figure 4A rows1 and 3**), the predictions at the number of cells/quadrats correctly classified show an improvement in the overall classification accuracy (upto 88.5%), especially for PD patients which improved from 66.7% to 69.39%, while also increasing the overall precision (**Appendix Figure 6 row3**). For the second crossover experiment, we cluster the cells across all quadrats from the pre-treatment cases (finer scale setting; see Methods), project the cells onto a 2D t-SNE, and depict the NN SDM disease predictions (generated using the coarse scale setting) to obtain a modest overlap and clearer distinction between PD (orange) and SD (green) regions (**Figure 5Ai, ii**). In the third crossover experiment, we use the coarser scale setting, where quadrats are sequential tiles of the images, and utilize the NN SDM predictions to color the quadrats for each FoV generating a risk map (**Figure 5B**). These risk maps are arranged in a 2D layout using Mistic^29^ where the 2D co-ordinates are generated by clustering the cellular marker densities for each FoV. Their orientation suggests that spatial measurements of the tumor ecology, and thus the interactions between markers, can be used to make accurate predictions of how a patient will respond to therapy. We observe that most of the PD images are colored in yellow/orange/red, correctly identifying these images, and therefore the patients, of having a higher chance to progress. Similarly, SD images have green/blue hues denoting a lower chance to progress. Additionally, we verified these differences using a statistical network model (**Appendix M.4**) that confirmed the presence of ecological diversity in pre-treatment biopsies from PD and SD patients.

**Figure 5:**
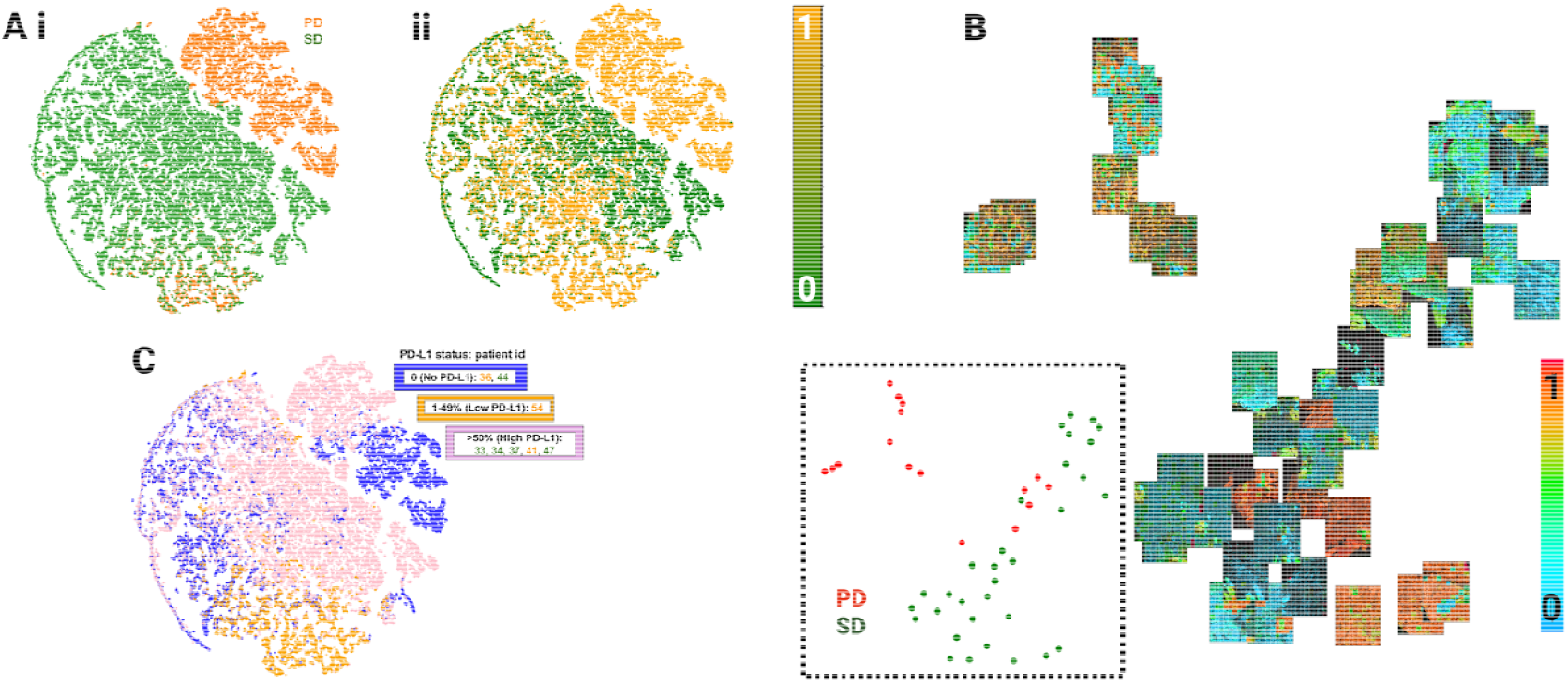
**A**. Combining cellular neighbors and quadrats using images from pre-treated patients where each quadrat is represented either as a cell with its neighbors (finer scale setting) or as a tile of 50um x 50um with groups of cells (coarser scale setting). In the former setting, we cluster the cells to identify cellular neighborhoods and plot the t-SNE showing **(i)** patient response, **(ii)** quadrat-trained NN SDM predictions with overall mean accuracy= 75%, mean accuracy for SD patients = 60% and for PD patients = 100%. **B**. In the coarse scale setting, we generate risk maps for each FoV where the quadrats are colored using the probability of progression scores and arranged into an image t-SNE using Mistic. Inset: original t-SNE showing disease response spread per FoV from pre-treatment. **C**. 2D t-SNE (similar to Figure 4Fi, shown in inset) depicting cells colored at the patient level and based on one of the three clinically annotated PD-L1 status categories: (0: No PD-L1; 1-49%: Low PD-L1; >50%: High PD-L1). Each patient is also clinically assigned to a response category: SD (green) or PD (orange) (inset).

The predictions from both the SVM and NN SDM are suggestive that distinct ecologies exist between PD and SD patients. This is consistent with prior studies that found infiltration of intra-tumoral CD3+ and CD8+ T cells in NSCLC is associated with better survival outcome^11,43,44^ and that FoxP3 cells are associated with a negative prognosis^45,46^. Further, in NSCLC patients, spatial proximity of tumor and FoxP3 cells has been found to be associated with diminished survival^45^. Additionally, since both the SVM and NN SDM leveraged the same predictive ecologies, it implies that these ecologies are inherently present in the data and are not artifacts of the prediction algorithms themselves.

### Potential of distinct ecologies as companion biomarkers for PD-L1 status

Through two tests, we further gauge the disease progression predictions using PD-L1 status alone, which is a recommended biomarker for NSCLC, particularly when deciding whether a patient is likely to benefit from immunotherapy treatments. In a first test, we train a SVM on the total number of cells (from cell segmentation) and a NN SDM on the total number of PPCs (from quadrats), using only the PD-L1 expression, from the pre-treatment images in FoVdata to predict patient response categories, i.e., SD versus PD (**Figure 4A rows 4-6)**. We show that using PD-L1 alone for response prediction has a predictive power of only 63% across both single cells and quadrats for FoVdata (**Figure 4A**).

In our second test, we utilize the clinically assigned PD-L1 status for each patient. The PD-L1 status is defined as the percentage of tumor cells that express PD-L1, and it is shown that tumors expressing high amounts of PD-L1 (50% or more) may respond particularly well to immunotherapy^47^. We rerender the FoVdata t-SNE in **Figure 5Ai** where cells per patient are colored according to the patient’s PD-L1 status (**Figure 5C)**. There are four clinically-assigned PD-L1 status categories for our nine-patient cohort: 0: No PD-L1; 1-49%: Low PD-L1; >50%: High PD-L1; Unknown: unable to assess PD-L1. Additionally, each patient is clinically categorized into either SD (green) or PD (orange) (**Figure 5A**). We observe that within each PD-L1 status category, there is a mix of both response categories. For instance, the No PD-L1 category has one PD and one SD patient, while the high PD-L1 category has four SD and one PD patient. Further, there was one patient in the unknown category that was assigned to be a PD patient. Note that for this patient, we only had one pre-treatment image and therefore the patient’s data was not used in the prediction experiments and is not depicted in **Figure 5C**. This shows that there were 3 misclassified patients out of 9, indicating that using PD-L1 status alone, the best patient response prediction we would achieve is 66.7%. In comparison, our ML predictions (**Figure 5Aii**) using PD-L1 in combination with other markers along with their spatial context, can predict patients with an accuracy between 75%-88%. This immediately highlights the clinical challenge of using PD-L1 status as a lone diagnostic biomarker as well as strengthens the utility of our marker combinations as a companion biomarker for PD-L1 status to predict treatment response.

## Discussion

Analyzing multiplexed images of patient tissues has the potential to become a routine clinical procedure for early detection, clinical cancer diagnosis, treatment planning and prognosis for many cancer types including NSCLC. We analyze multiplexed images to quantify the spatial ecology of tumor and immune cell types to understand if these ecologies can predict disease progression dynamics under therapy. Through two cornerstone image processing techniques (cell segmentation and image tiling to generate quadrat counts) combined with spatial statistics, machine learning, and deep learning, we perform in-depth analyses of multiplexed images based on the frequency, phenotype, and spatial distribution of immune and tumor cells within the immune landscape of NSCLC. Multiplexed field of views (FoVs) were obtained from pre- and during-treatment (days 15-21) biopsies in nine patients with advanced/immunotherapy-refractory NSCLC treated with an oral HDAC inhibitor (vorinostat) combined with a PD-1 inhibitor (pembrolizumab). We characterize cellular composition and spatial neighborhoods that predict distinct tissue ecologies in patients whose NSCLC has progressive disease or stable disease following treatment. We have identified potential biomarkers for treatment response, e.g., PD-1+PanCK+FoxP3. We demonstrate that PD tumors are characterized by a highly suppressed immune response (due to presence of FoxP3+ and PD-1+ immune cells) prior to treatment, whilst SD tumors showed a higher colocalization of PD-L1+CD3+CD8+ with tumor cells. Together, these findings indicate that there are distinct cellular compositions, both within the tumor regions and at the tumor boundaries, for PD versus SD patients. Furthermore, we use these distinct biomarkers to train support vector machines (SVMs) and neural network species distribution models (NN SDM) to predict disease progression.

Previous studies have shown that HDACi increases anti-PD-1 response by enhancing tumor immunogenicity, in part through upregulation of major histocompatibility complex (MHC) and T-cell chemokine expression^48^. As HDACi alone is not effective in NSCLC^47^, we hypothesize the TME associations described here are primarily associated with response to anti-PD-1. In this study, we have shown that both single cell and quadrat analysis methods can be synergized to predict disease progression, as well as visualize these predictions at the patient level through risk maps (**Figure 5B)**. Further, these methods are generalizable across various cancer types as well as imaging modalities. We identify that disease progression predictions made by SVMs using PD-L1 alone have an overall mean accuracy of 63% (when using both cell segments and quadrats (at the patient level, **Figure 4A**; at the cell/quadrat level per patient, **Appendix Figure 6**)). On the other hand, through our approach of using multiple markers along with PD-L1, the overall mean accuracy using SVM trained on cells is 88%, the NN SDM trained on quadrats is 75%, and the weighted SVM combining both cells and quadrats is 88.5% (**Appendix Figure 6**).

The predictions obtained from single cells are generally better than and comparable to those from the quadrats. This is mainly due to the marker resolution, i.e., in single cells, each marker is analyzed at the cellular level whereas in the quadrats, each marker represents a group of cells present within that quadrat. For research questions related to cellular co-localization and its impact on cellular processes (such as, cell signaling), one can rely on cellular analysis approaches. For studies investigating global interactions of cells (such as, interaction networks), quadrat approaches are appropriate. Nevertheless, it is important to note that the predictions in this study, generated by utilizing all the markers along with their spatial context, are notably higher than those obtained by using PD-L1 alone (**Figure 4A**).

Through these findings, our study reinforces that there are technical and clinical challenges in using PD-L1 as a standard diagnostic biomarker^5–8^. Thus, there is a need for complementary diagnostic biomarkers along with PD-L1 for different cancer types and treatment^28^. From both our single-cell and quadrat analyses, we find the tumor tissue immune architecture in pre-treatment biopsies is different in responding and non-responding tumors. Herein, we have shown that these pre-treatment ecologies of NSCLC patients are distinct, so much so that their spatial organization (CNs or QNs) can be used to predict response to treatment more accurately than PD-L1 alone. In PD patients, pre-treatment biopsies demonstrated exhausted T cells (PD-1) and immune suppressive Tregs (FoxP3), along with decreased CD8 T-cell infiltration. However, in SD patients, we observe increased CD8 infiltration in tumor regions. These findings are in alignment with studies that show PD tumors to exhibit more immune suppressed environments, and SD tumors to have higher numbers of infiltrated immune cells^49^. Additionally, we verified these differences using a statistical model (**Appendix M.4**) that confirmed the ecological diversity in pre-treatment biopsies from PD and SD patients.

The current study does have its limitations. One limitation is that it uses retrospective clinical trial data. In addition, it has a small sample size (FoVdata: n = 9 patients, 92 FoVs) with stable and progressive disease cases, but no complete or partial response cases. While selection bias may be present in FoVdata, we minimized this risk by selecting FoVs that had a tumor to stromal compartment ratio close to 0.5. Also, our potentially significant clinical observation - that the pre-treatment tissue ecology is predictive of response or resistance to a treatment - is in the context of combination therapy, i.e., with HDAC inhibitor combined with a PD-1 inhibitor. In another ongoing study, we compare these predictive ecologies with a monotherapy cohort. Our major motivations to embark the current study on this limited cohort were: a) to glean insights into the presence and effect of spatiotemporal ecologies in shaping treatment resistance as very little is known about the impact of drugs on evolving ecologies in NSCLC; b) to demonstrate the technical feasibility of our combined image analyses methodology; and c) as a computationally-driven study following the promising clinical observations inferred from FoVdata^28^ where such predictive biomarkers were hypothesized to exist.

Acknowledging these limitations, we are currently expanding this study to validate the biomarkers from this initial study in a larger cohort of patients, including those from the ongoing randomized Phase II clinical trial (NCT02638090). This larger cohort has patients with a wider range of responses, including stable and progressive disease, complete responders, and partial responders. Additionally, this trial has two arms, one in which anti-PD-1 naïve NSCLC patients are treated with either anti-PD-1 pembrolizumab monotherapy (Arm A, n=39), or with pembrolizumab + HDACi vorinostat (Arm B, n=39; similar to FoVdata in this study). Given the complexity of HDACi effect on the TME as well as uncertainty of eventual approval of HDACi treatment of NSCLC patients in combination with PD-1 inhibitors, our future work will be aimed at better understanding how the biomarkers might vary in a monotherapy versus combination therapy setting as well as across patient response categories. This will help interpret the variations in marker interaction and tumor architecture unique to each therapy. Further, using Mistic and our statistical network analysis to study varying networks across treatment time points, we will investigate marker changes that characterize immune-cool tumors from immune-hot tumors. We will also develop mechanistic mathematical models to simulate response to treatment, which will aid in clarifying *why* the markers are predictive.

In summary, we use single cells and quadrats combined with spatial statistics, machine learning, and deep learning, to perform in-depth analyses of multiplexed images (FoVs) based on the frequency, phenotype, and spatial distribution of immune and tumor cells within the immune landscape of advanced/immunotherapy-refractory NSCLC. We demonstrate that PD tumors are characterized by a highly suppressed immune response (due to presence of FoxP3+ and PD-1+ immune cells) prior to treatment, whilst SD tumors showed a higher colocalization of PD-L1+CD3+CD8+ with tumor cells. This is consistent with previous literature that in NSCLC patients, spatial proximity of tumor and FoxP3 cells results in exhausted immune cells and has been associated with diminished survival^45^, while infiltration of CD3+ and CD8+ T cells along with anti-PD-L1 treatment is associated with better overall survival^11,43,44,49^. These, therefore, validate our cell- and quadrat-based approaches, and further indicate that our dual approaches are applicable beyond NSCLC as well as the specific subset of markers discussed here, and will be even more relevant for use with new markers where the underlying biology is not clear.

## Methods

### Non-spatial analyses

To compare the underlying patterns of tumor evolution in NSCLC between PD and SD patients, we examined the density of markers across the FoVs (FoVdata dataset) using basic ecological analysis, for example, average quadrat counts. We view each FoV as a collection of fixed-size patches, called quadrats, defined as partitions of an image into multicellular regions, creating sequential tiles in order. Each quadrat is 100µm x 100µm in size and expresses the area normalized sum of marker positivity (after thresholding) of all cells present within that quadrat. In **Figure 2A**, an example 7-marker FoV (with tumor regions in cyan) is shown (**i**) along with its corresponding cell segments (**ii**) and quadrats (**iii**). We plot the marker abundances (based on average positive pixel counts (PPC)) across all the quadrats of PD and SD patients, before and during treatment (**Figure 2B**). We further investigate the difference in PD and SD marker densities by looking at the non-metric multidimensional scaling (NMDS)^50^ ordination of the positive pixel counts (PPCs) for each FoV. NMDS shows the tumor composition of each treatment category. This is then followed by a confirmatory permutational analysis of variance (PERMANOVA^51^) test on the ecological dissimilarity matrices (computed using PPCs of the quadrats) (**Figure 2C**). Additionally, using the PPCs, we plot the pairwise marker correlations across SD and PD patients (**Figure 2D**).

### Spatial analyses with cell segments

Using our multiplexed imaging pipeline for single cells (**Appendix Figure 1i, Methods M.1.1, M.2**), we extract cell segments and tumor regions per FoV. Once the cell segments are extracted using a U-Net^52^, we build a count matrix with cells as rows and markers as columns, and cluster the count matrix to identify heterogeneous cell types using a Gaussian mixture model (GMM)^30^. To understand the spatial organization of tumor and immune cells across PD and SD patients, we investigate each cell in the context of its spatial neighbors. This is done by generating cellular neighborhoods (CNs) where the marker expression of each cell is the average of ten of its nearest spatial neighbors in Euclidean space (**Figure 3A-C**). A CN can be defined as the minimal set of cell types that are both functionally and spatially similar. This approach is inspired by a previous CN implementation^53^ with the major difference that we do not have a fixed set of cell types at the onset but work with cell types that are organically inferred through the clustering process. Along with the cell segments, we also computationally demarcate tumor-rich regions (approximated using multiple convex hulls; **Appendix M.1.2)**, across images, that are higher in PanCK expression (red regions in **Appendix Figure 1ii**, right panel). To compute the colocalization of markers, we use a modified Morisita-Horn co-localization index^16,54^ (**Appendix M.5**).

### Spatial analyses with quadrats

In addition to the cell segments, we process image quadrats^55^ (**Appendix M.1.3**) by identifying quadrat neighborhoods (QNs) (**Figure 3D, E**) and building species association networks^31–33^ (SANs) (**Figure 3F**). Similar to CNs, QNs are defined as the minimal set of functionally and spatially similar quadrats. These are extracted using the Leiden clustering algorithm^36^ so as to identify spatial communities. SANs provide a modelling framework to extract direct spatial associations based on spatial co-localization, while removing indirect effects. This is a popular approach in quantitative ecology used by landscape ecologists^56,57^. The fundamental modeling steps in SANs are data preparation, model fitting and assessment, and prediction. The quadrats are combined and fit with a Gaussian copula graphical lasso (GCGL) using the *ecocopula* package^34,35^ applied to pre- and during-treatment images.

### Approaches to combine cell segments and quadrats

We describe three approaches to combine the insights derived from cell segments and quadrats. In the first crossover experiment, we created a weighted cell marker matrix where each cell’s marker expression is multiplied by the average of its corresponding quadrat progression probabilities (as obtained from the NN SDM). This weighted marker matrix is then used to train the SVM. The motivation to use such a weighted approach was to primarily handle unbalanced sample sizes (5 SD and 3 PD patients in FoVdata), as well as unbalanced regions of disease progression within each FoV. We observe that this approach leads to an improvement in the overall classification accuracy at the cell/quadrat level per patient, especially for PD patients (**Appendix Figure 6**), and that these results can be further augmented by additional patient samples.

For the second approach, we describe a crossover experiment between the cell segmentation and quadrat approaches to further reinforce the distinct architectures existing between PD and SD patients (**Figure 5A,B**). For this, we use images from only the pre-treatment cases and redefine the quadrat into both finer and coarser scales, and cluster the quadrats. In the finer scale, each quadrat is represented as a cell with its neighborhoods (**Figure 5Ai, ii**), and in the coarser scale, each quadrat is a fixed-size patch with a collection of cells and their neighborhoods (**Figure 5B**).

In our third approach to quantify those markers that potentially drive the patients from the pre-treatment state to the on-treatment state, we build a statistical model to infer the difference in marker interactions, in the form of a network, between these patient categories. This is done by utilizing the SANs (**Figure 3F**) from the FoVs, across response groups and across pre- and during-treatment states. The inference problem can be stated as a least-squares minimization problem^58^ and can be further regularized using prior biological knowledge of marker interactions (**Appendix M.4, Appendix Figure 5**).

### Support Vector Machine (SVM) used to make predictions with single cell data

We implemented the SVM classifier using Python’s scikit learn’s^59,60^ (sklearn) *SVM, SVC* classes. The classifier uses a linear kernel. Classification statistics (accuracy, precision, recall, and F1-score) were calculated using scikit learn’s *metric* package. The LOOCV was implemented using the *LeaveOneOut* class provided in sklearn’s *model_selection* suite.

### Neural Network Species Distribution Model (NN SDM) for predicting disease progression using quadrats

We use a feed forward neural network^61^ (NN) to identify species (marker) distribution interactions (SDM) to predict disease progression in patients (**Appendix Figure 7**). The inputs to the NN SDM are quadrats generated from the FoVdata images. Quadrat dimensions for the NN SDM are 50μm x 50μm. The activation function used is Leaky Rectified Linear Unit^62^ (ReLU). The loss function used is cross balanced loss^63^, which was minimized using the AdamW^64^. Due to the unbalanced nature of the data, the Synthetic Minority Over-sampling Technique (SMOTE)^65^ was used to oversample the minority class, thereby balancing the training data. To analyze feature importance (**Figure 4E**), we use integrated gradients^41^ available through PyTorch’s model interpretability library, Captum^42^.

### Training classifiers for predicting disease progression

Using FoVdata, we trained and tested classifiers based on support vector machines (SVMs), and neural network species distribution networks (NN SDMs) to predict disease progression of patients. The response of the patients was known *a priori*. Using the Leave-one-out cross-validation (LOOCV) setup, at the patient level, the classifiers were trained on the training set and predictions of disease progression were made on the test set. To generate these predictions, we used images from 8 patients (5 SD and 3 PD) from the 9-patient cohort since one patient (classified as PD) had only a single image taken before treatment.

We used both SVMs and NN SDMs as these are popular and well established supervised learning techniques to build prediction models^66^. While SVMs are powerful for finding optimal decision boundaries, especially in high-dimensional spaces, and with non-linear separation boundaries, NN SDMs are considered non-parametric, making no assumption on the distribution of data and the structure of the true data generator. The similarity of biomarkers obtained by both the SVMs and NN SDMs show that the results obtained are not artefacts of the specific prediction algorithm used but are prominent underlying patterns in the data.

## Data availability

The NSCLC images reported in this study cannot be deposited in a public repository because they are unpublished and shall be made available upon reasonable user request. To request access, contact lead contact, Alexander R. A. Anderson. Patients were assigned to SD/PD groups based on their response and RECIST criteria. No cell lines were used in this study. All analyses were performed blinded such that experimenters performing data analysis were unaware of the patients.

As an example dataset, we have deposited an anonymized subset of 10 FoVs from the NSCLC dataset here: https://doi.org/10.5281/zenodo.6131933.

## Code availability

Code pipeline for the cell segmentation-based analysis and quadrat-based analysis will be made available upon user request. Mistic is downloadable at: https://github.com/MathOnco/Mistic.

## Acknowledgment

The authors gratefully acknowledge funding by the National Cancer Institute via the Cancer Systems Biology Consortium (CSBC) U01CA232382 and U54CA274507 (supporting S.P., C.G., M.R.-T., R.A.G., A.R.A.A), and support from the Moffitt Center of Excellence for Evolutionary Therapy (supporting M.R.-T., R.A.G., and A.R.A.A).

Figures are created using Inkscape, BioRender.com, NN-SVG^67^ and Keynote.

## Author contributions

Conceptualization, S.P., C.G., M.R.-T., and A.R.A.A.; methodology, S.P, C.G., M.R.-T., and A.R.A.A.; writing – original draft, S.P., C.G., M.R.-T., A.B., J.G., S.A., R.A.G., and A.R.A.A.; writing – review and editing, S.P., C.G., M.R.-T., A.B., J.G., S.A., R.A.G., and A.R.A.A.; software design and implementation, S.P. and C.G.; formal analysis, S.P. and C.G.; validation, S.P., C.G., and A.R.A.A.; software testing, S.P. and C.G.; supervision, S.A., R.A.G., and A.R.A.A.; funding acquisition, S.A., R.A.G., and A.R.A.A.

