## Supplementary methods and figures for "Distinct tumor-immune ecologies in NSCLC patients predict progression and define a clinical biomarker of therapy response"

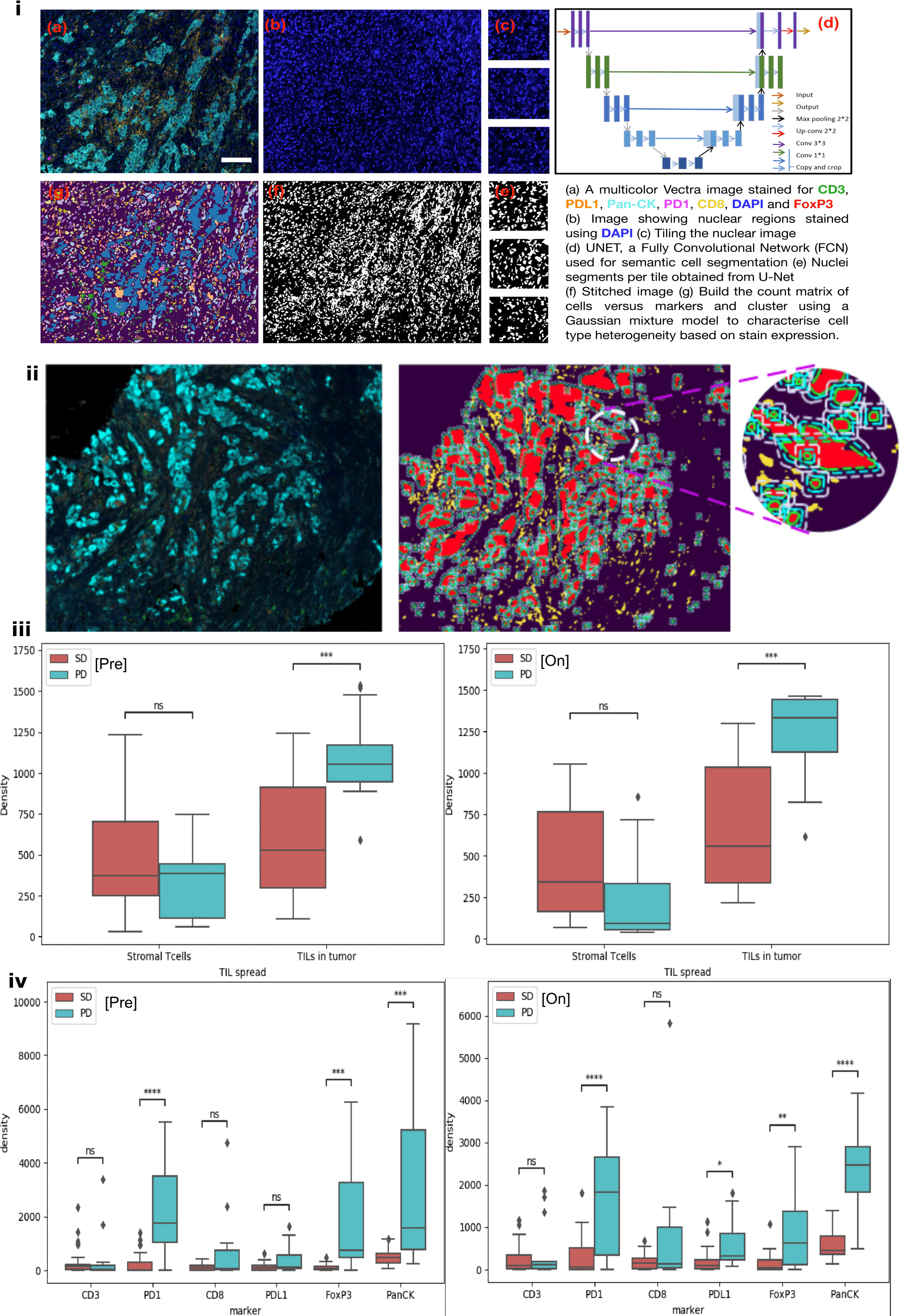

**Appendix Figure 1**:

**(i) Image processing pipeline:** (a) the input is an original 7-stain Vectra image with DAPI (blue; nuclei), CD3 (green; T cells), CD8 (yellow; effector T cells), FoxP3 (red; regulatory T cells), PD-1 (pink; inhibitory receptor on immune cells), PD-L1 (orange; immune checkpoint marker), and pan-cytokeratin (PanCK, cyan; epithelium) (b) the image with corresponding DAPI stains showing nuclei is extracted. The nuclear-stained image is tiled (c) and fed into a deep-learning architecture called U-Net (d) to identify cell segments. (e) U-Net identified cell segments per tile which are then stitched to create the image in (f). A count matrix with cells (rows) and markers (columns) is computed from (f) and is used for clustering to identify heterogeneous cell types. In (g) cell segments are colored based on the cluster assignments obtained through a Gaussian mixture model. Scale bar = 200 pixels/mm.

**Quantifying the tumor-immune cells at the tumor border using convex hull approximation:**

**(ii)** (**Left**) An example Original 7-color Vectra image multiplex FoV, (**Middle**) Tumor regions (in red) are identified by extracting PanCK-rich regions from cell segmentation and the tumor region boundaries are approximated using convex hulls. Multiple convex hulls are drawn extending into and outside the tumor region to aid quantifying the immune-tumor cells colocalized at the tumor boundary. Inset: Zoomed in region showing tumor regions in red and three convex hull boundaries in dotted lines.

**(iii)** (**Left**) Box plots showing the presence of Stromal immune cells and immune cells within tumors in SD and PD patients in pre-treatment and (**Right**) on-treatment cases. For both pre and on treated cases there is a higher presence of immune cells in tumor in PD than SD patients. The Bonferroni-adjusted p-value with significance are: *: 1.00e-02 < p <= 5.00e-02, **: 1.00e-03 < p <= 1.00e-02, ***: 1.00e-04 < p <= 1.00e-03, ****: p <= 1.00e-04.

**(iv)** (**Left**) Box plots showing number of immune and tumor cells in the tumor border in SD and PD patients pre-treatment and (**Right**) on-treatment cases. For both pre and on treated patients there is a higher presence of immune cells (PD-1, FoxP3) with tumor (PanCK) in PD than SD patients. The p-values are Bonferroni-adjusted p-values with significance as in (iii).

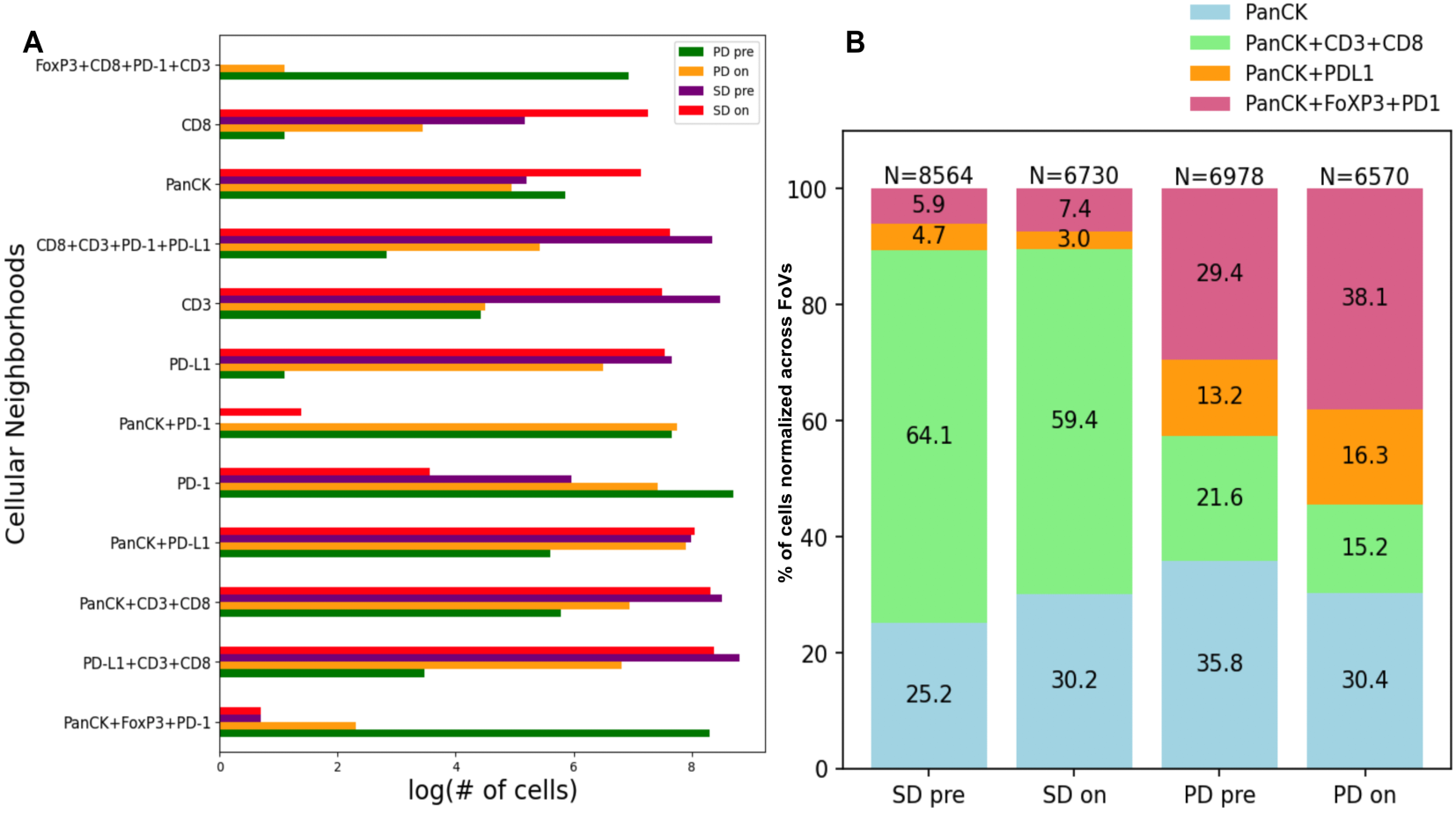

**Appendix Figure 2:**

**A.** Log normalized number of cells across various spatial cellular neighborhoods and four response categories (PD pre, PD on, SD pre, SD on).

**B.** Relative proportions of tumor cells (PanCK+), and tumor cells colocalized with one or more of the functional markers (CD3+ T cells, CD8+T cells, FoxP3+ (Treg cells), PD-L1, and PD-1) within each patient response category based on treatment timing. Pooled data from all patients are shown with cell numbers per category shown above the respective bar. Colocalization is calculated based on the weighted Morisita-Horn measure (Appendix Method M.4)

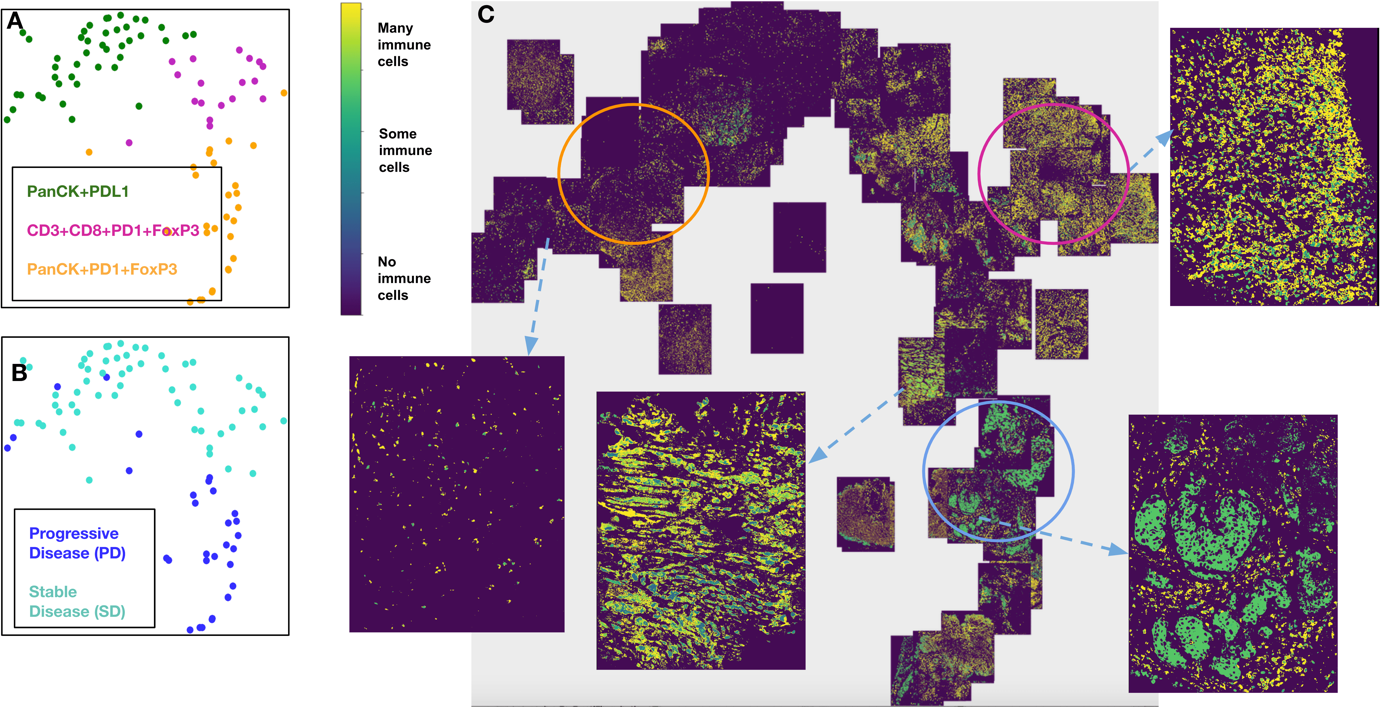

**Appendix Figure 3**: PD and SD patients have significantly different cellular compositions.

**A.** 2D t-SNE plot showing three clusters annotated with the differentially expressed markers per cluster. Clusters are obtained using the tumor-immune cell counts at the tumor border approximated by convex hulls (see **Figure 2**).

**B.** Same t-SNE as in **(A)** depicting the disease response spread.

**C.** Image t-SNE using same t-SNE coordinates as in **(A)** illustrating the gradient of immune cells (CD3, CD8, PD-1 and FoxP3) and tumor cells (PanCK) across images. A higher colocalization of PanCK+PD-1+FoxP3+ (shown in green) is seen for PD patients. Circles denote regions of immunologically cold (orange), immunologically warm (blue) and immunologically hot (pink) tumors.

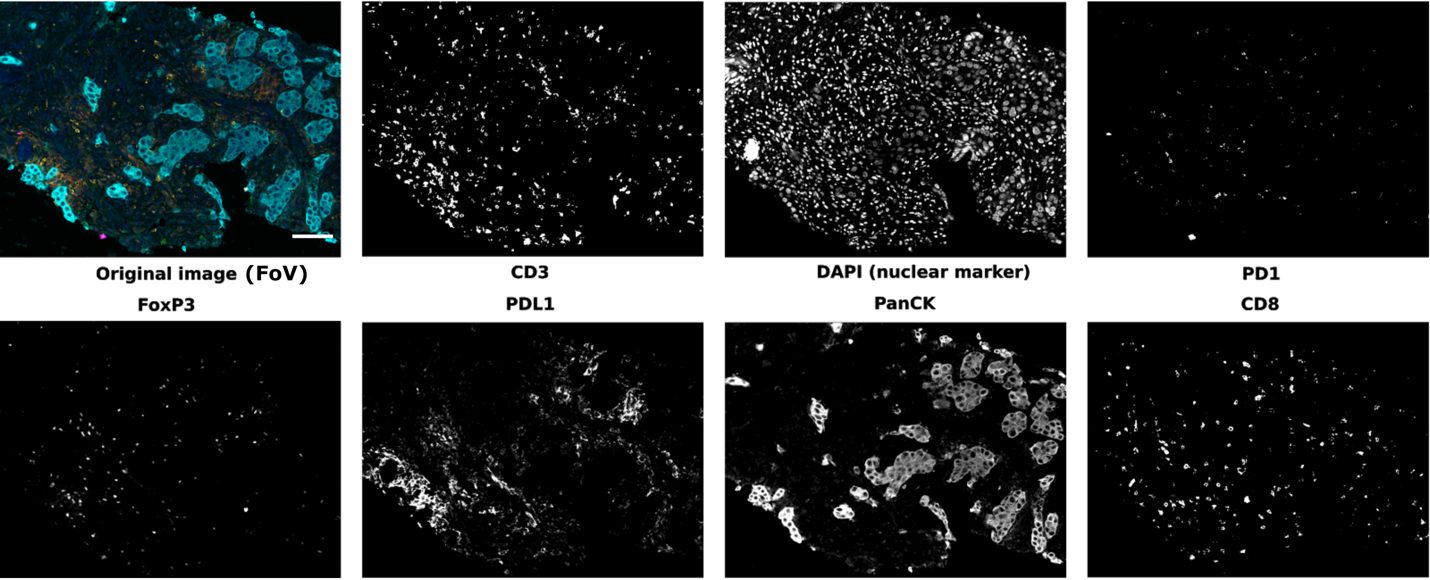

**Appendix Figure 4**: Example marker panels for a FoV from the 9-patient NSCLC cohort (FoVdata). Scale bar = 200 pixels/mm.

The markers for the 7-stain Vectra image (FoV) are DAPI (nuclei), CD3 (T cells), CD8 (effector T cells), FoxP3 (regulatory T cells), PD-1 (inhibitory receptor on immune cells), PD-L1 (immune checkpoint marker), and pan-cytokeratin (PanCK; epithelium).

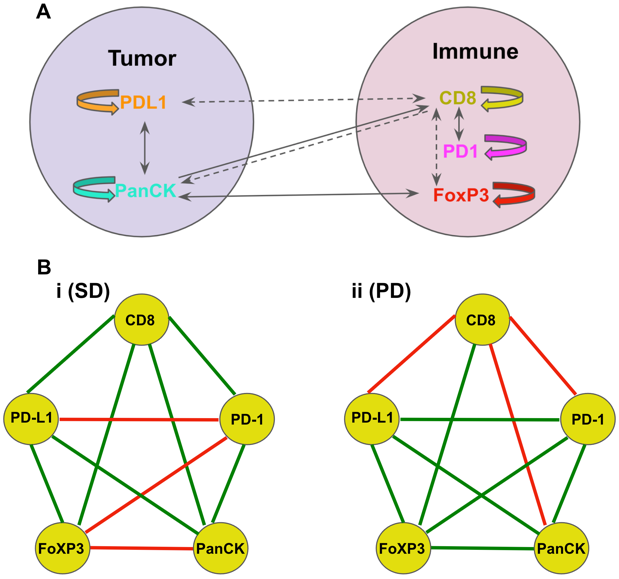

**Appendix Figure 5:**

**A.** The prior biological interactions between tumor and immune markers used in the *difference network* analysis. These interactions are encoded in a matrix, $W^{prior}$, as: +1 (solid links: positive interaction); -1 (dotted links: negative interaction); and 0 (no interactions)

**B.** $W^{LSprior}$ depicted as a network for SD (i) and PD (ii) obtained by solving the constrained least-squares optimization problem in Eqn 4. $W^{LSprior}$maps the markers from the ’pre’ to the ’on’ state while incorporating known mechanisms as prior information. Green links indicate inferred positive marker associations and red lines indicate inferred negative associations.

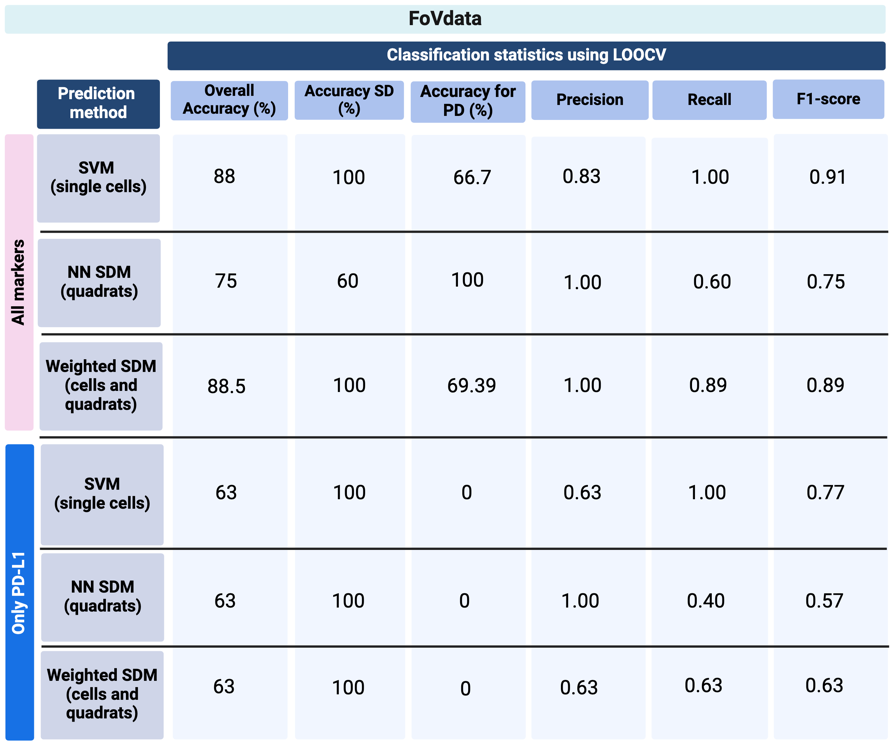

**Appendix Figure 6:** Disease progression prediction metrics using a Support Vector Machine (SVM) trained on cells, neural network species distribution model (NN SDM) trained on quadrats, and weighted SVM trained using quadrat probability distributions, from pre-treatment FoVdata images. Metrics shown for all markers (analyzed in their spatial context) versus PD-L1 alone. Metrics calculated based on the number of cells/quadrats correctly classified, per patient.

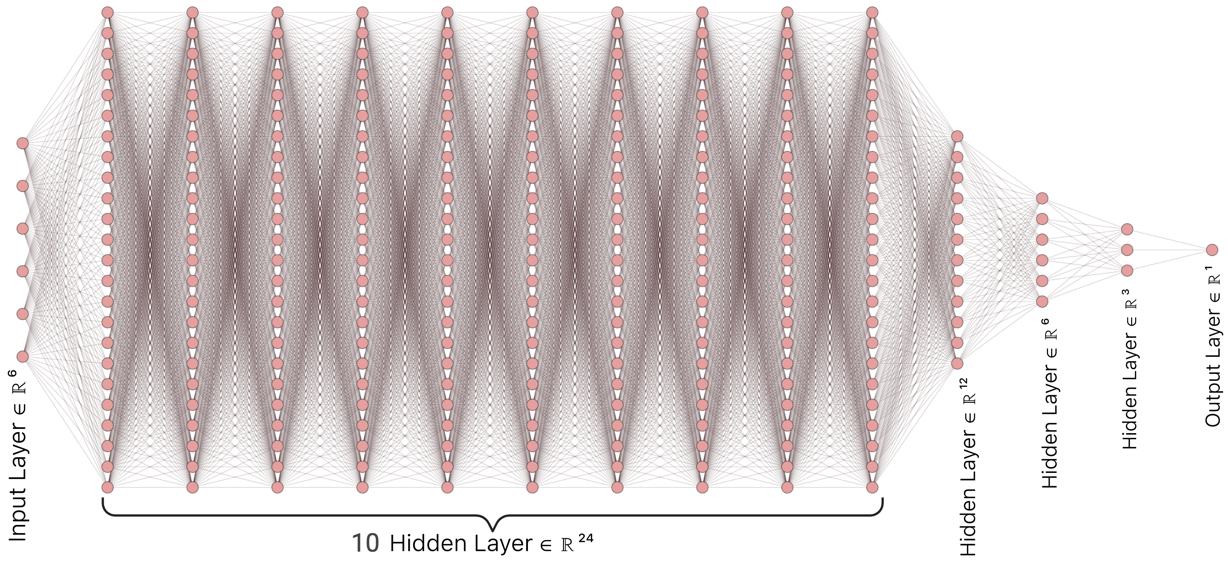

**Appendix Figure 7:** NN SDM architecture with one input layer, 10 hidden layers, 3 funnel layers, and one output prediction layer.

### **Methods**

M.1.1 Multiplexed image processing pipeline for cell segmentation in FoVs

Each 7-stain Vectra multiplexed image is available as a tiff stack with 7 sub tiffs: one tiff for each marker. We have built an image processing pipeline that consists of three parts to analyze each tiff stack (**Appendix** **Figure 1i**):

(1) the pre-processing unit that denoises and cleans each 1008-pixel x 1344-pixel sub tiff of the 7-stain Vectra Tiff stack (**Appendix** **Figure 1ia**). We start with generating the grayscale image of each sub tiff and choose the nuclear image stained with DAPI for cell segmentation (**Appendix** **Figure 1ib**). Since the images are ridden with technical noise, it is crucial to extract true stain signal from noise. For this we apply an Otsu thresholding to the images which involves iterating through all the possible threshold values and calculating a measure of spread for the pixel levels each side of the threshold, i.e., the pixels that either fall in foreground or background. Foreground pixels are regarded as true stain signals while background pixels are noise. Next, the images are tiled into 256-pixel x 256-pixel frames (**Appendix** **Figure 1ic**).

(2) The cell segmentation unit: The image tiles are used as input to U-Net^1^, a Fully Convolutional Network, for cell segmentation (**Appendix** **Figure 1id**). We further trained U-Net using nuclear segments available from Kaggle’s Data Science Bowl.

(3) Downstream analysis unit: Once the cell segmentation predictions are made per tile (**Appendix** **Figure 1ie**), we stitch the tiles (**Appendix** **Figure 1if**) and perform clustering on the stitched image (**Appendix** **Figure 1ig**). For clustering, we use a Gaussian Mixture model^14^ which identifies heterogeneous cell types based on presence of stain expression, and does not require the number of clusters to be known in advance.

In the FoVs, cell segments consisted of cells from tumor and stromal regions.

M.1.2 Convex-hull approximation

To demarcate tumor regions within an FoV, we draw convex hulls around those clusters (obtained from the above cell segmentation and clustering process) that are high in PanCK. For this we use the *scipy.spatial.ConvexHull*^6^ function that finds the smallest convex set that encloses all the region coordinates, forming a convex polygon. The coordinates for plotting the polygon are obtained from the *regionprops*^7^ function generated during cell segmentation. Once the initial convex hulls are identified, the process is repeated to plot hulls extending both inside and outside the initial hulls using fixed boundary offsets. For example, a fixed offset id is ‘added’ to the coordinates of the initial hulls to plot hulls extending ‘outside’ the initial hulls.

M.1.3 Multiplexed image processing pipeline for quadrat counts

The quadrat-based analyses are based on marker abundances in the entire quadrat (i.e., those possibly outside nuclear masks), and so used a slightly different approach to remove channel bleed and noise.

(1) Markers were assigned into mutually exclusive groups, i.e., those expressed only on T-cells (CD3, CD8, FoxP3, PD-1), or tumor cells (PanCK, PD-L1). Next, each pixel in the image is assigned to one of those groups based on which marker had the maximum expression. For example, if a pixel has fluorescence for PanCK and FoxP3, it would be assigned to the T-cell group if FoxP3 (or any other T-cell marker) is greater than the expression of PanCK (or any other tumor marker), or tumor group if PanCK is higher than FoxP3. Once assigned to a group, the pixel values in the other group's channel(s) are set to zero. The result is that a pixel can be positive for more than 1 marker in the same group (i.e., T-cell that is CD3+/CD8+), but not for markers in mutually exclusive groups (i.e. not possible to have CD3+/PanCK+).
(2) Channel bleed is further reduced by subtracting the intensity of the adjacent channel, but only if that other channel belongs to a mutually exclusive group and uses an adjacent fluorophore. For example, if a pixel is positive for FoxP3 (Opal 650) and PanCK (Opal 690), and FoxP3 expression is greater than PanCK, then the pixel is classified as T-cell and the PanCK channel is subtracted from the FoxP3 channel.
(3) After removing channel bleed, each channel is thresholded to determine the number of pixels that are positive for each marker. In the case of the FOV, Otsu thresholding was used to separate foreground (pixels positive for each marker) from background).

M.2 PD patients have high PD-1 and T regulatory cells (FoxP3+) in both the tumor border and tumor regions

Using our multiplexed imaging pipeline for single cells (**Appendix Figure 1i**), we extract the cell segments per FoV using a U-Net^1^, build a count matrix with cells as rows and markers as columns, and cluster the count matrix to identify heterogeneous cell types (refer **Appendix Methods M.1.1**). From these cell types, we automatically demarcate tumor-rich regions, across images, that are higher in PanCK expression (red regions in **Appendix** **Figure 1ii**, right panel). The tumor regions are approximated using multiple convex hulls to allow for spatial quantification of tumor and immune cells within and outside the tumor border.  We note that for both pre and on treated patients, there is a higher presence of immune cells within the tumor (**Appendix** **Figure 1iii**) and a higher abundance of tumor and immune cells across the tumor border (**Appendix** **Figure 1iv**) in PD than SD patients signaling that there are distinct cellular compositions for PD and SD patients.

M.3 Quantifying the tumor-immune cell colocalization at the tumor border

Next, to quantify the tumor-immune cell colocalization at the tumor border we cluster the tumor-immune cell counts at the tumor border approximated using convex hulls (see **Appendix Figure 1ii,** right panel). Clustering is performed using a Gaussian mixture model^8^ (GMM). The input matrix to the GMM consists of cells (as rows) and their marker distribution as features (columns). The rows of the input matrix are ordered based on the cluster assignments and the z-score of the marker expression (columns) is averaged over vectors per cluster representing a cell type (rows). Those markers that have a higher z-score per cluster are identified as the differentially expressed markers for that cluster. The clusters are visualized using a 2D t-SNE plot where each point abstracts an image (**Appendix** **Figure 3A, B**). We obtain distinct clusters showing higher colocalization of PanCK+PD-1+FoxP3+ for PD patients and higher colocalization of PanCK+PD-L1+ along with CD3+CD8+ immune cells for SD patients indicating underlying structural differences between PD and SD patient groups. Further, to better understand the relationship of the clusters to the actual images, we generate an image t-SNE (**Appendix** **Figure 3C**) where we replaced each dot in the t-SNE of **Appendix Figure 3A, B** with the corresponding multiplexed images. This image t-SNE was rendered using Mistic^9^ and this arrangement of images as an image t-SNE clearly highlights the difference in immune cell abundance across PD and SD patient groups. The image t-SNE also helps in identifying groups of immunologically hot, warm or cold tumors across patients based on immune cell colocalization at the tumor margins and tumor regions.

M.4 Inferring underlying networks of SD and PD patients using prior information

To quantify the distinct architectures across PD and SD patients, we build a statistical model to infer the difference in marker interactions, in the form of a network, between these patient categories. This is done by: utilizing the species association networks^10–12^ (SANs; Figure 3F) along with the edge weights, and marker expression both at the pre-treatment and on-treatment states for each category; and capturing the ‘*difference network’* that potentially drives the patients from the pre-treatment state to the on-treatment state. The inference problem can be stated as a least-squares minimization problem^8^ and can be further regularized using prior biological knowledge of marker interactions.

Problem formulation

We denote the change in the i^th^ marker expression in the pre-treatment case at time *t*, m_i_^pre^(t), as:

Equation 1:

$\frac{dm_{i}^{pre}\left( t \right)}{dt}= \sum_{j=1}^{n} W_{ij}m_{j}^{pre}\left( t \right)- m_{i}^{on}\left( t \right)$

where *n* is the number of markers, *i, j* = 1, 2, …, *n*, and *W* is the weight matrix. Equation 1 can be written in matrix form as:

Equation 2:

$$\frac{dM^{pre}\left( t \right)}{dt}= WM^{pre}\left( t \right)- M^{on}\left( t \right)$$

where $M^{*}\left( t \right)=m_{1}^{*}\left( t \right), m_{2}^{*}\left( t \right), \ldots,m_{n}^{*}\left( t \right)$ and * denotes pre or on states. We can solve Equation 2 as a least-squares minimization problem given *n* markers and ($M_{i}^{pre}, M_{i}^{on}$) as steady states. This can be written as:

Equation 3:

$$W^{LS}=arg\min_{W} f\left( W \right)= arg\min_{W} \sum_{j=1}^{n} ||WM^{pre}- M^{on} ||$$

Further, we can add prior knowledge of biological mechanisms between markers as additional constraints to the least-squares problem **(Appendix Figure 5A)**. These mechanisms can be captured in pairwise format as a matrix $W^{prior}$ where a 1 indicates positive interaction, 0 indicates no interaction and -1 indicates negative interaction. Equation 3 can be written as follows to yield $W^{LSprior}$:

Equation 4:

$$W^{LSprior}=arg\min_{W} f\left( W \right)= arg \min_{W} \sum_{j=1}^{n} ||WM^{pre}- M^{on} ||+\lambda||W o W^{prior} ||$$

Implementation

We construct weighted count matrices for the four categories: SD (pre, on) and PD (pre, on), by combining the SANs and the edge weights obtained in **Figure 3F**. Starting with the SAN for each category, a weighted adjacency matrix, $A^{adj}$, is created. $A^{adj}$ is a square distance matrix representing connections between markers, wherein ‘1’ indicates a marker-marker connection, and ‘0’ otherwise, and each connection is multiplied by weights from the SAN where the weights are bound by -1 (red/negative association) and 1 (blue/positive association). For each $A^{adj}$, we construct a similarity matrix, $S^{adj}$, using$S^{adj}={-0.5(QA}^{adj}Q^{t})$ with $Q_{ij}=\delta_{ij}-1/n$ where $\delta$ is the Kronecker delta function and $n$ is the number of markers, based on the centering operation in kernel PCA^8^. The weighted count matrix, $X^{adj}$ can be computed by the eigenvalue decomposition^8^ of $S^{adj}$. Assuming that each marker follows a Gaussian distribution, we can model these weighted count matrices of cells by markers to follow a multivariate Gaussian distribution, without loss of generality. We compute the first and second order moments to describe each of the four categories. Next, for each category, we randomly draw 10,000 samples creating a random matrix of 10000 rows and 5 markers. This is repeated 100 times and the random matrices are averaged out to create one representative matrix denoting that category. This is done to ensure that we work with random matrices that a) are invariant to biases and shifts while generating them from distance and similarity matrices and b) have the same number of rows in the ’pre’ and ’on’ states that obey the observed/empirical distributional moments. Implementing Equation 4 by incorporating the biological prior, gives us the optimized $W^{LSprior}$ for SD and PD (**Appendix Figure 5B i,ii**) respectively. It becomes evident through $W^{LSprior}$ for SD (**Appendix Figure 5Bi**) that there is an increased CD8 interaction with the tumor, a decrease in FoxP3 interaction with tumor and a positive interaction between CD8, PD-L1 and PanCK. For PD (**Appendix Figure 5Bii**), the $W^{LSprior}$ indicates an increase in FoxP3 interaction with tumor, a decrease in CD8 interaction with tumor and a positive interaction between FoxP3, PD-1 and PanCK. Note that these interactions that differentiate SD from PD were previously identified from the image t-SNE analysis (see **Appendix Figure 3**) as well as the quadrat analysis using (**Figure 3D-F**).

M.5 Weighted Morisita-Horn measure

To quantify the co-localization of tumor and immune cells regions, we use the Morisita-Horn (MH) similarity measure^13,14^. For two populations, immune and tumor, with proportions $p_{imm}$ and $p_{tum}$ respectively, the MH measure can be written as: $\frac{2{* p}_{imm}{* p}_{tum}}{p_{imm}2+ p_{tum}2}$.

We consider a weighted form of the MH measure where weights across cells of the two populations are calculated using inverse-distance weighting. This weighting influences the similarity measure based on the cells that are closer to, than farther away from, the cell in question. The weighted MH measure is given as:

$$\frac{2(\sum_{imm \epsilon I} \frac{p_{imm}}{d_{avg}(tum, imm)})*\sum_{tum \epsilon T} \frac{p_{tum}}{d_{avg}(tum, imm)})}{{(\frac{p_{imm}}{d_{avg}(tum, imm)}}^{2}+{\frac{p_{tum}}{d_{avg}(tum, imm)}}^{2})}$$

where $d_{avg}(tum, imm)$ is the average distance of a tumor cell, *tum*, to its neighboring immune cell, *imm*. *I* is the total number of immune cells and *T* is the total number of tumor cells.

### **List of abbreviations**

DAPI: 4′,6-diamidino-2-phenylindole

CD3 (T cells): cluster of differentiation 3

CD8 (effector T cells): cluster of differentiation 8

CN: cellular neighborhood

FoVs: field of views (FoVs)

FoxP3 (regulatory T cells): forkhead box P3

GCGL: Gaussian copula graphical lasso

GMM: Gaussian Mixture model

HDAC: Histone deacetylases

MHC: major histocompatibility complex

NMDS: non-metric multidimensional scaling

NSCLC: Non-Small Cell Lung Cancer

PanCK: Pancytokeratin

PERMANOVA: permutational multivariate ANOVA

PD: progressive disease

PD-1 (inhibitory receptor on immune cells): Programmed cell death protein 1

PD-L1 (immune checkpoint marker): Programmed death-ligand 1

PPC: positive pixel counts

QN: quadrat neighborhood

SAN: species association network

SD: stable disease

SVM: support vector machines

TME: tumor microenvironment

t-SNE: t-distributed stochastic neighbor embedding

UMAP: Uniform Manifold Approximation and Projection
